# Inhalation of marijuana affects *Drosophila* heart function

**DOI:** 10.1101/459792

**Authors:** IM Gómez, MA Rodríguez, M Santalla, G Kassis, JE Colman Lerner, O Aranda, D Sedán, D Andrinolo, CA Valverde, P Ferrero

## Abstract

Medical uses of marijuana have been recently approved in many countries, and after a long ban on research, there is despicable scientific evidence regarding its action and side effects. We investigated the effect of inhalation of vaporized marijuana on cardiac function in *Drosophila melanogaster*, a suitable genetic model for assessing cardiovascular function. Chronic exposure of adult flies to vaporized marijuana reduces heart rate, increments contractility and prolongs relaxation. These changes are manifested in the cardiomyocytes with no effect in calcium handling, and in the absence of the canonical cannabinoids receptors identified in mammals. Our results are the first evidence of the *in vivo* impact of phytocannabinoids in *D. melanogaster* and open new paths for genetic screenings using vaporized compounds, providing a simple and affordable platform prior to mammalian models.

## Introduction

Human body normally produces endocannabinoids, molecules that trigger receptor-mediated signaling pathways (1). The endocannabinoid system regulates a variety of physiological processes such as release of neurotransmitters, perception of pain and cardiovascular, gastrointestinal and hepatic functions. The pharmacological manipulation of endocannabinoid levels or the administration of cannabinoid agonists such as those from *Cannabis sativa* has been used for the treatment of various pathologies. The marijuana plant produces more than 60 terpenophenolic cannabinoid compounds in its trichomes, present in leaves and flowers. The main cannabinoids are cannabidiol (CBD), tetrahydrocannabinol (THC) and cannabinol (CBN). A meta-analysis of cannabinoids for medical purposes provided evidence on their efficacy in pain control, antiemetic in chemotherapy, stimulation of appetite in patients with HIV, modulation of sleep disorders, motor dysfunction in paraplegia, anxiety disorders, diabetes and metabolic syndrome, among others (2). However, the therapeutic potential of marijuana, as well as the possible collateral effects of its use, has not been fully exploited due to the long lasting ban of its use. The treatments of patients with marijuana extracts or cannabis oil, although being illegal in many countries, continued in an intuitive manner. The recent lift of the ban for medical and even recreational use of marijuana in a handful of countries shall commit the scientific community to build a body of knowledge and establish animal models suitable for basic research and translational medicine. In this respect, the study of side effects must be carefully evaluated. For instance, the acute effects of cannabinoids in the cardiovascular system results in the increase of heart rate and vasodilatation, although hemodynamics effects are attenuated with repeated doses. Smoking or inhaling marijuana has been associated with increased risk of infarct and angina in patients with heart diseases (3). However, we still lack an exhaustive analysis of cardiac performance after exposure to cannabinoids. Results arising from animal models are complex, according to experimental conditions and route of administration, being the inhalation pathways the least explored. Herein, we study the effect of vaporization of marijuana and its cannabinoids in the cardiac performance of *Drosophila melanogaster* a rising experimental model for cardiac function.

## Results

Canonical receptors for endocannabinoids and phytocannabinoids such as those present in mammals are absent in *Drosophila* genome (4, 5). However, the fruit fly possess its own endogenous cannabinoids (6) thereby we hypothesized that this organism may be responsive to phytocannabinoids.

To test this hypothesis, we developed a system for administration vaporized substances to the intact *Drosophila* flies. The marijuana’s extract utilized was provided by the Argentinian NGO “Mamá Cultiva”, that it is currently used in the treatment of patients with autism. We first determined the proportion of THC, CBD and CBN by gas chromatography and mass spectrometry (Table S1). The marijuana’s strain used in the present study is enriched with THC (114:1 THC/CBD). Flies were exposed to extract of *Cannabis sp*. vaporized at 188°C (370° F) using a commercial vaporizer (Da Vinci) connected to the device shown in Fig. 1A. At this temperature, most cannabinoids were present in the vapor (Table S2). By controlling a manifold stopcock, vapor was suctioned and transferred by a syringe into a vial containing the flies to be treated. The compounds were incorporated through and distributed by the tracheal ramifications of the flies. Although it is a passive process, we defined it as inhalation. Two groups of flies were exposed for two different periods (6-8 days and 11-13 days) and compared with flies not exposed to marijuana (control). A third group (T&I) received vaporized cannabinoids for five days and then deprived of cannabinoids for the following five days. Fig. S2 shows accumulative amount of cannabinoids vaporized during the treatment period to which each group of flies was exposed. According to the World Health Organization, a human treated chronically is one whose treatment is applied for one or more years. For a human average lifespan of 74 years, this represents > 1.4 % of lifespan expectancy. In accordance to this definition, and considering that *Drosophila melanogaster* has a lifespan expectancy of 40 days at 29°C, the cannabinoid treatment applied for 6-8 or 11-13 days, represent 13% or 30% of their average lifespan, respectively (Fig. 1B). Therefore, the conditions implemented in this study are to be considered a chronic exposition of flies to cannabinoids. The chronic exposure does not significantly affect lifespan (Fig. 1C), therefore, it allows us to assess the cardiac activity in a normal viable population.

**Figure 1.**
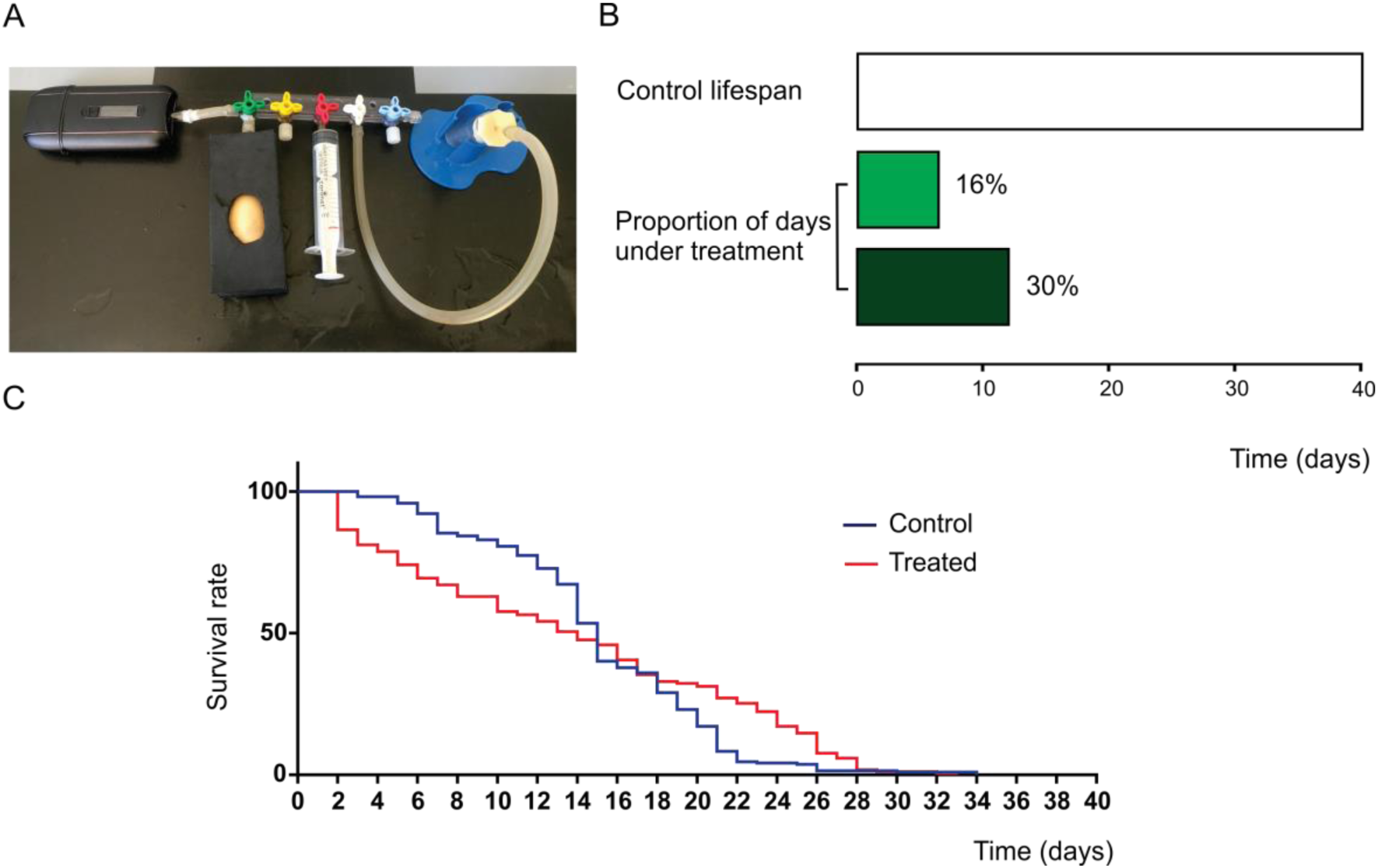
Chronic exposure of *Drosophila melanogaster* to vaporized extract of *Cannabis sp* does not affect lifespan. A. Device connected to a vaporizer designed for providing vaporized cannabinoids into a vial containing the group of flies subjected to the cannabis treatment. B. Representation of the lifespan percentage of flies submitted to cannabinoids exposition. C. Kaplan-Meier survival analysis for treated (red) and not treated (blue) adult flies with extract of *Cannabis sp*. Cumulative survival is depicted over time. Both curves are not statistically significantly different according to the log-rank (Mantel-Cox) test. Control: N= 217 Treated: N=170.

Cardiac function was assessed in semi-intact heart preparations as we have previously escribed (7, 8). A reporter system that express the green fluorescent protein (GFP) in cardiac and pericardial cells was used to track the movement of a cardiac or pericardial cell during each cardiac cycle (systole and diastole). Mechanical parameters were defined as previously described (9). Fig. 2A shows representative mechanical recordings of cardiac activity of the four groups of flies (control, 6-8 days treated, 11-13 treated, T&I). Inhaled marijuana reduced heart rate (negative chronotropic effect) as a function of the exposure time (Fig. 2B). This reduction was significantly different from control in individuals with the longer exposure time to the drug. Interestingly, heart rate reduction was significantly higher in flies in which treatment was interrupted for 5 days after 5 days of treatment (indicated as T & I in Fig. 2B). This result indicates that a shorter time of exposure is enough to induce heart rate reduction, although the appearance of the effect was delayed. The mechanism of the delayed effect suggests either an indirect cardiac response mediated by a neurohormonal pathway activity or a non-immediate processes of gene expression regulation triggered in the cardiac tissue that manifests after stimulus was interrupted.

**Figure 2.**
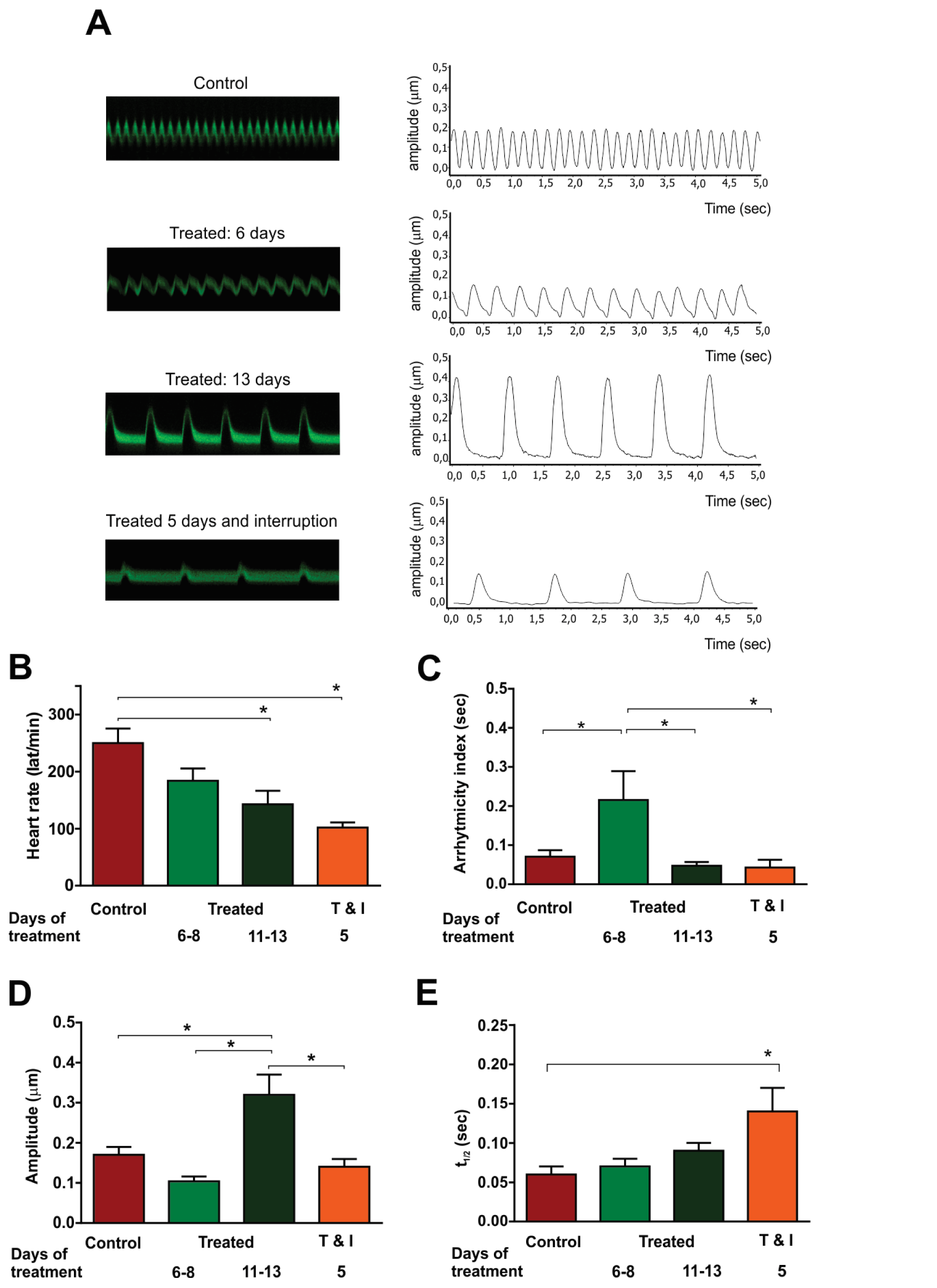
Cardiac contractiliy is modified by extract of *Cannabis sp*. A. Typical recordings (left) and digitalized signals of heart wall displacement for each group during 5 sec are shown. Variation of heart rate and contractility are evident. B. Average results indicate heart rate reduction in treated group of flies. N = 14, 9, 7, 10, Control, treated 5-8 days, 10-13 days and 5 days and interruption (T&I) respectively.C. Arrhytmicyty Index is increased in the group submitted to the shorter period of exposition to the extract. N = 14, 9, 7, 10. D. Average results of wall displacement. Longer period of exposition to the extract (10-13 days) significantly enhances contractility. N = 20, 9, 7, 12. E. Average results of half relaxation time. Prolongation of time of relaxation is induced by 5-8 days of exposition but evidenced later during adult life. N = 14, 9, 7, 12.

Exposure to the cannabinoids extract induced an increase in Arrhytmicity index (Fig. 2C), in individuals treated during 6-8 days, which would indicate that this phenomenon is an immediate and transitory effect since it was not observed either in T & I individuals or in individuals exposed to the drug during 11-13 days.

We then explored the effect of different inhalation time on cardiac contractility evaluated by means of wall displacement. Fig. 2D shows that wall shortening increased significantly after prolonged inhalation of *Cannabis sp* extracts (11-13 days). Although at first sight this positive inotropic response may be related to the slowdown of heart rate (9), contractility in the T&I group is similar to the control, in spite that heart rate remains reduced. Moreover, we previously observed that increasing heart rate reduces Ca^2+^ transient with minimal effects on wall shortening (9), an opposite pattern to the one described here for cannabinoids. Therefore, these results would suggest that the increase in contractility observed in the 11-13 treated flies might be mediated by mechanism different from heart rate reduction. Heart relaxation estimated by half relaxation time (t½) shows a trend to increase (negative lusitropic effect) with the time of exposure to vaporized extract. Although we can argue that this trend to increase is associated with the associated decrease in heart rate, the difference in relaxation time of treated versus non-treated flies only attained statistically values in the T&I group (Fig. 2E). This might suggests again that this prolongation of relaxation may be associated to non-immediate mechanisms.

Aging reduces heart rate and prolongs relaxation in humans and *Drosophila melanogaster* (7, 8, 10). In order to dissect a possible incidence of aging on heart rate reduction and relaxation prolongation, observed in the T&I group, we then compared aged-matched flies. The results obtained confirm that both parameters were significantly modified after cannabinoids exposure (Fig. S2). Similar effects on contractility were observed with the treatment (Fig. S3).

Calcium signaling is essential for cardiac contraction (systole) and relaxation (diastole) that occur when electrical activity is followed by a mechanical action in the cardiomyocytes. The link between the electrical and mechanical activity is denominated Excitation-Contraction Coupling (ECC), which depends on the intracellular Ca^2+^ concentration (Ca^2+^ transient). After an action potential depolarized the cell membrane, Ca^2+^ increases in the cytosol and binds to the myofilaments to induce its contraction. For relaxation to occur, Ca^2+^ is quickly removed from the cytosol mainly by the sarcoplasmic reticulum (SR) Ca^2+^ATPase pump (SERCA) and the Na^+^/ Ca^2+^ exchanger (NCX). Slow removal systems like mitochondrial Ca^2+^ transporter and sarcolemmal Ca^2+^ATPase pump (PMCA) contribute to a lesser extent to reduction of Ca^2+^_i_ (11). These mechanisms govern cardiac contractility in *Drosophila* and mammals (12). Therefore we studied whether mismanagement of Ca^2+^ is involved in the cardiac phenotype induced by the exposure to marijuana.

We exposed to vaporized cannabinoids 10 days old flies bearing a genetically-encoded Ca^2+^ reporter (G*Ca*MP3) in the heart. Cytosolic Ca^2+^ transients were measured and SR Ca^2+^ content was estimated by application of a 10 mM caffeine pulse applied on the semi-intact heart preparation (Fig. 3A). This pharmacological intervention, widely used in mammal studies, opens the sarcoplasmic reticulum Ca^2+^ channels (ryanodine receptors, RyR) allowing SR- Ca^2+^ release. Amplitude of caffeine-induced Ca^2+^ transient estimates the SR- Ca^2+^ content. Comparison of pre- caffeine and caffeine Ca^2+^ transients decay, allows to calculate the relative importance of each rapid removal system during relaxation. We determined that Ca^2+^ transient, SR Ca^2+^ content as well as the activity of the main proteins involved in Ca^2+^ handling were indistinguishable in control flies when compared to treated flies (Fig. 3B-F). Therefore we concluded that Ca^2+^ handling is not modified after cannabinoids treatment in *Drosophila*, and that the positive inotropic effect exhibited in the treated group might be due to increased myofilament Ca^2+^ sensitivity (Fig. 2D and Fig. S3). Further studies are needed to investigate the mechanism of this effect.

**Figure 3.**
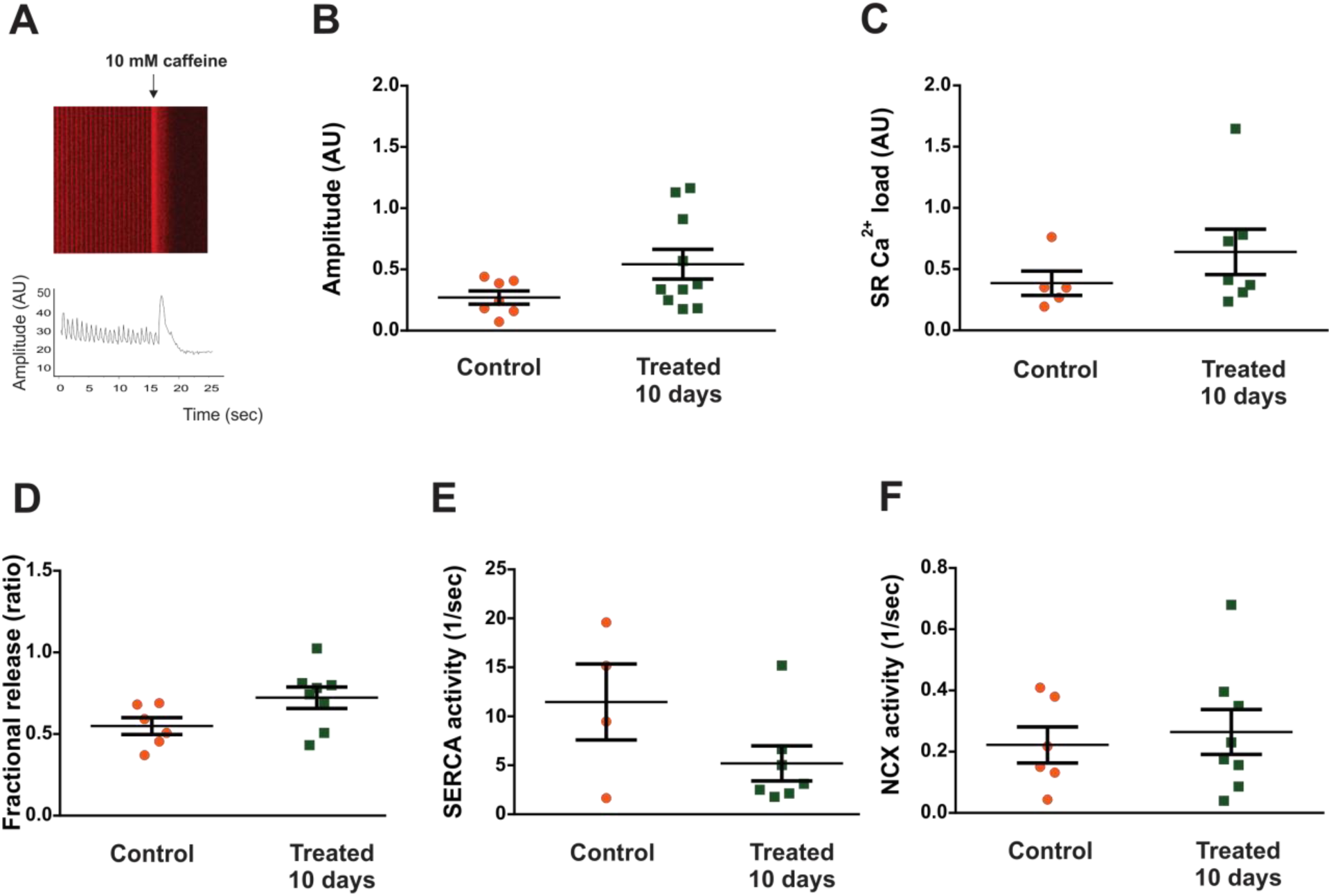
*Ca*lcium management is not modified by cannabinoids in *Drosophila* heart. A. Image and representative tracing of Ca^2+^ transient. A pulse of 10 mM caffeine is applied to the semi-intact preparation and visualized as a sudden increase in the fluorescent signal. Amplitude of caffeine-induced Ca^2+^ transient estimates the SR Ca^2+^ content. Fractional release indicates activity of RyR because it represents the fraction of SR Ca^2+^ that is released during each systole referred to SR Ca^2+^ content. On the other hand, decay of caffeine induced ^*^Ca2+^*^ transient could be attributed mainly to NCX activity and the pre-caffeine Ca^2+^ transient decay results from the combined SERCA and NCX activity. All parameters are similar between both groups of flies compared (control and exposed during 10 days to extract of *Cannabis sp*) B: Ca^2+^ transient (C: N = 7, T: N = 10) C: SR Ca^2+^ load (C: N = 5, T: N = 7). D: Fractional release that indicates RyR activity (C: N = 6, T: N = 8). E and F: SERCA and NCX activity, respectively (C: N =4, T: N =7; C: N = 6, T: N = 8).

Altogether, effects of chronic exposure on heart performance can be categorized on the following: 1- early immediately visualized changes like heart rate variability, 2- early changes that become evident later in the life of flies as reduced heart rate and prolongation of relaxation, and 3- changes produced by prolonged exposure to cannabinoids like the increment in *Drosophila* heart contractility.

## Discussion

Inhalation of vaporized marijuana is an important route to supply cannabinoids to patients for pain control in arthritis, multiple sclerosis and some types of cancer (13). It is faster than oral delivery for cannabis components to the blood torrent. However, the study of vaporized cannabinoids in animal models are difficult to set up out due to the costs of animal facilities maintenance and the complex logistic of vaporization in large volumes. We developed a system for providing vaporized substances to numerous individuals in small volumes and for studying chronic exposure in a short period of time on account of the relative short life cycle of flies. This method has been successfully extended to other disease related models such as tobacco (14) opening the way to large screenings at affordable costs.

One contraindication for medical use of cannabinoids is their impact on heart function. Smoking cannabis might cause angina in patients with heart disease and increments the risk of infarct in patients who already had myocardial infarction (3). However, endocannabinoids and synthetic cannabinoids seem to prevent ischemic damage in rat models, mainly through CB2 type endocannabinoid receptors (15-17). However, several aspects remain unsolved: actions of marijuana’s extract, effects of chronic administration, detailed cardiac information from healthy individuals, and the impact of inhalation route. The cardiac phenotype in healthy adult flies provoked by chronic exposure to *Cannabis sp* reduces the heart rate as occurs with repeated exposure to cannabinoids in humans (18). In *Drosophila*, this could be attributed to intrinsic functions of the heart, as the fruit fly has an open vascular system (19). Like humans, *Drosophila* heart exhibits calcium L and T type calcium channels (11). It is feasible that cannabinoids can affect these voltage operated channels, as well as it was described for neurons in mammal models where THC reduces T type calcium channel (20). Our analysis indicates that SR calcium handling is not affected and similar results were observed in rat isolated myocytes exposed to the endocannabinoide anandamide (21).

We cannot rule out the possibility that cannabinoids modulate neural functions because of its impact on heart rate. For example, the endocannabinoid anandamide inhibits adrenaline release from peripheral sympathetic nerves, inducing vasodilatation (22). Moreover, in mammals, cannabinoids inhibits glutamate release (19) and glutamate receptors are expressed in human heart (20). It might be possible that cannabinoids modulate adrenergic and glutamatergic input on cardiac tissue. The presence of octopaminergic – a beta adrenergic like pathway - and glutamatergic terminals that innervate the *Drosophila* heart (23, 24) opens the possibility that cannabinoids might modulate heart activity through these pathways. It has being shown that application of glutamate to *Drosophila* heart accelerates the cardiac action potential frequency (23). Therefore, reduction of glutamatergic signaling might modulate cardiac activity slowing heart rate, under cannabinoids influence.

Endocannabinoids have been described for *Drosophila*, (6) however the canonical receptors could not be identified in the genome. It has been proposed that cannabinoid receptors evolved from the last common ancestor of bilaterians, with a secondary loss occurring in insects and other clades. The evolution of cannabinoid receptors in invertebrates has been controversial and evidence organized in hierarchical analysis levels: *in vivo*, in vitro or in silico. The absence of canonical receptors in insects has been inferred from in vitro and in silico studies (5), however we provide here the first systemic response to vaporized cannabinoids in the cardiac function in *Drosophila*. These findings and the method we set up render the fruit fly a useful platform for low cost and low- to high-throughput phytocannabinoids induced cardiac phenotypes screenings and new target discovery.

## Materials and Methods

### *Drosophila* stocks, rearing and crosses

Fly stocks were amplified and maintained in vials at 28°C, partially filled with a mixture of maize flour, glucose, agar and yeast supplemented with 10% antimycotic. Assays were made with the wild type strain *Ca*nton-S (BDSC# 9514) (25) obtained from Bloomington *Drosophila* Stock Center. The handC-GFP line, expressing the reporter protein Green Fluorescent Protein (GFP) in the heart and pericardial cells was used for contractility assessment. For cytosolic calcium sensing, we used flies harboring the reporter system G*Ca*MP3 specifically expressed in heart cells (UAS-G*Ca*MP3/UAS-G*Ca*MP3; tinCΔ4-Gal4, UAS-G*Ca*MP3/tinCΔ4-Gal4, UAS-G*Ca*MP3) (26). HandC-GFP flies were provided by Prof. Achim Paululat, and G*Ca*MP3 expressing flies were provided by Dr Heiko Harten, both from University of Osnabrück, Germany (27).

WT flies were breeded with individuals carrying one of these reporters. We analyzed heterozygous individuals of F1 that possess one copy of the reporter system in order to obtain information on mechanical activity (using the handC-GFP reporter) or Ca^2+^ transients (using the UAS-G*Ca*MP3 reporter).

### Extract characterization

For chemical characterization vegetal material was extracted with ethanol. An aliquot of the ethanolic extract was subjected to a clean up treatment with C18, activated carbon and MgSO4 to adsorb impurities and contaminants. Clean extract was diluted in hexane and injected in a PerkinElmer Clarus 650 gaseous chromatograph, coupled to a PerkinlElmer Clarus SQ 8 S mass spectrometer. Determination of THC, CBD and CBN was conducted following the technic described by Tayyab and Shahwar (2015) (28). Analytical standards of THC, CBD and CBN employed were purchased from Cerilliant Corporation. Concentrations of THC, CBD and CBN were determined (expressed in mg of cannabinoid / g of vegetal material) in the *Ca*nnabis strain utilized for the experiments.

### Administration by vaporizing extracted cannabinoids

A costumed device was connected through one end to an Ascent Portable Vaporizer (Da Vinci) containing 0.03 g of vegetal extract. Temperature was set to 370°F (187.78°C), for vaporizing all types of cannabinoids present in the extract. The produced vapor was collected by a syringe, and 40 cc of vapor was insufflated in a 15.5 ml plastic vial containing adult flies. The top end of the vial was adapted with a filter for avoiding increase in air pressure within the vial when vapor was insufflated. Vaporized individuals received two daily doses and remained in contact with the vaporized substances for 15 minutes each dose before being transferred to their regular cultivation vials.

### Survival analysis

Populations of synchronized flies were obtained and separated in two groups. The first group received vaporized substances twice every day. The protocol was continued until the last individual died. The second group did not receive vaporized extract and was treated as control flies. Survival analysis was carried out counting live and dead flies every day. Statistical comparison and curves were obtained using the Kaplan-Meier method and analyzed using the logrank (Mantel-Cox) test.

### Evaluation of cardiac function in semi-intact heart preparation

The dissection of adult hearts was performed as described by Santalla et al. (2014) (7). The procedure was performed on a Schonfeld Optik model XTD 217 stereomicroscope. Individuals (7 and 11 days old flies) were briefly anesthetized with carbon dioxide (CO_2_), and placed in a 60 mm Petri dish coated with a thin layer of petroleum jelly and fixed by the dorsal region. The head and thorax were removed in this preparation; thus neuronal influence on the cardiac activity was not significant. The middle ventral region of the abdomen was opened and the internal organs were removed. The preparation was submerged in oxygenated artificial hemolymph solution containing 5 mM KCl, 8 mM MgCl_2_, 2 mM *Ca*Cl_2_, 108 mM NaCl, 1 mM NaH_2_PO_4_, 5 mM HEPES,4 mM NaHCO_3_, 10 mM trehalose, 10 mM sucrose, pH 7.1. *Ca*lcium recordings and mechanical activity of these semi-intact heart preparations were carried out using a *Ca*rl Zeiss LSM410 and LSM800 confocal microscopes.

### Mechanical activity and calcium transient recording

Mechanical activity was recorded from all flies harboring the GFP protein under the control of the handC driver, expressed in cardiomyocytes and pericardial cells. A pericardial or a cardiac cell was focused with a 20X objective. We then tracked the fluorescent signal of a cardiac or pericardial cell edge from one side of the heart. We verified that the movement of the pericardial or cardiac cells is synchronic and that they displace to the same extend. The movement of a single cell expressing the GFP reporter was followed by laser scanning in line scan mode by setting a line longitudinally to the displacement of either a pericardial cell or a cardiomyocyte. Recordings were obtained for 5 seconds. Cytosolic Ca^2+^ was assessed by the fluorescence produced by ^*^Ca2+^*^ binding to the G*Ca*MP3 reporter. Semi-intact preparations were visualized using a 5X objective and the laser was focused to stimulate a minimal central region of the conical chamber where the intensity of signal was the highest. Recordings of 60 sec allowed us to obtain a pattern of Ca^2+^ transients.

The images obtained were processed with a custom-made algorithm for Anaconda developed in our laboratory for obtaining over a threshold value, the intensity of fluorescence from a pericardial/cardiac cell. The images were obtained sequentially in time and then converted into a digitalized image of cell displacement. After conversion of the image to a txt file containing fluorescence values and time, the files were analyzed with LabChart software (AD Instruments, CO, USA).

We measured the displacement of the heart wall, expressed in µm from the diastolic to the systolic positions of a cardiac or pericardial cell. We measured the heart period as the interval between two consecutive contraction peaks, and then we calculated the heart rate. Arrhytmicity index (AI) was calculated as the standard deviation of periods normalized by their average. We also evaluated the time to half relaxation (t1/2), defined as the time from maximum contraction until half of the maximum relaxation (minimum contraction). For Ca^2+^ transient signal, we measured fluorescence intensity [(Fmax-F0)/F0)]. For estimating SR- Ca^2+^ load, fractional release, SERCA and NCX activities, a pulse of 10 mM caffeine was applied to semi-intact heart preparation of G*Ca*MP3 flies by means of a micropipette. Amplitude of caffeine-induced Ca^2+^ transient represents an estimation of SR- Ca^2+^ load meanwhile fractional release results from the ratio between pre-caffeine and caffeine-induced Ca^2+^ transient amplitudes. NCX activity was estimated as the inverse of the relaxation time constant (tau) of the caffeine-induced Ca^2+^ transient. SERCA activity was evaluated as the difference between the inverse of tau from pre-caffeine and caffeine-induced Ca^2+^ transients (1/tau pre-caff – 1/tau caff).

### Statistical analysis

One-way analysis of variance (ANOVA) followed by Tukey's post hoc test, was utilized for comparison of the differences among three or more groups. Student's t-test was used to evaluate differences between two groups. A p-value < 0.05 was considered significant.

## Acknowledgements

We want to thanks to Ana Clara Maldonado for her technical assistant and the non-governmental organization “Mamá Cultiva” for providing vegetal material utilized in our assays. We also want to offer our recognition to Dr. Marcelo Morante for helping us to generate a framework to develop research in our country. We thank Dr. Alicia Mattiazzi and Dr Rolando Rivera Pomar for their critical revision of the manuscript.

## Author contributions

Investigation and formal analysis: for chemical characterization: IMG, DA, DS, EC, OA: for physiological analysis: IMG, MAR, MS, PF. Conceptualization, project administration, supervision: PF. Methodology: CV, GK, PF. Writing – review & editing: CV, PF. Resources: PF.

## Data and materials availability

All data is available in the main text or the supplementary materials.

## Supplementary Materials

Figures S1-S3

Tables S1-S2

---

**Fig. S1.**
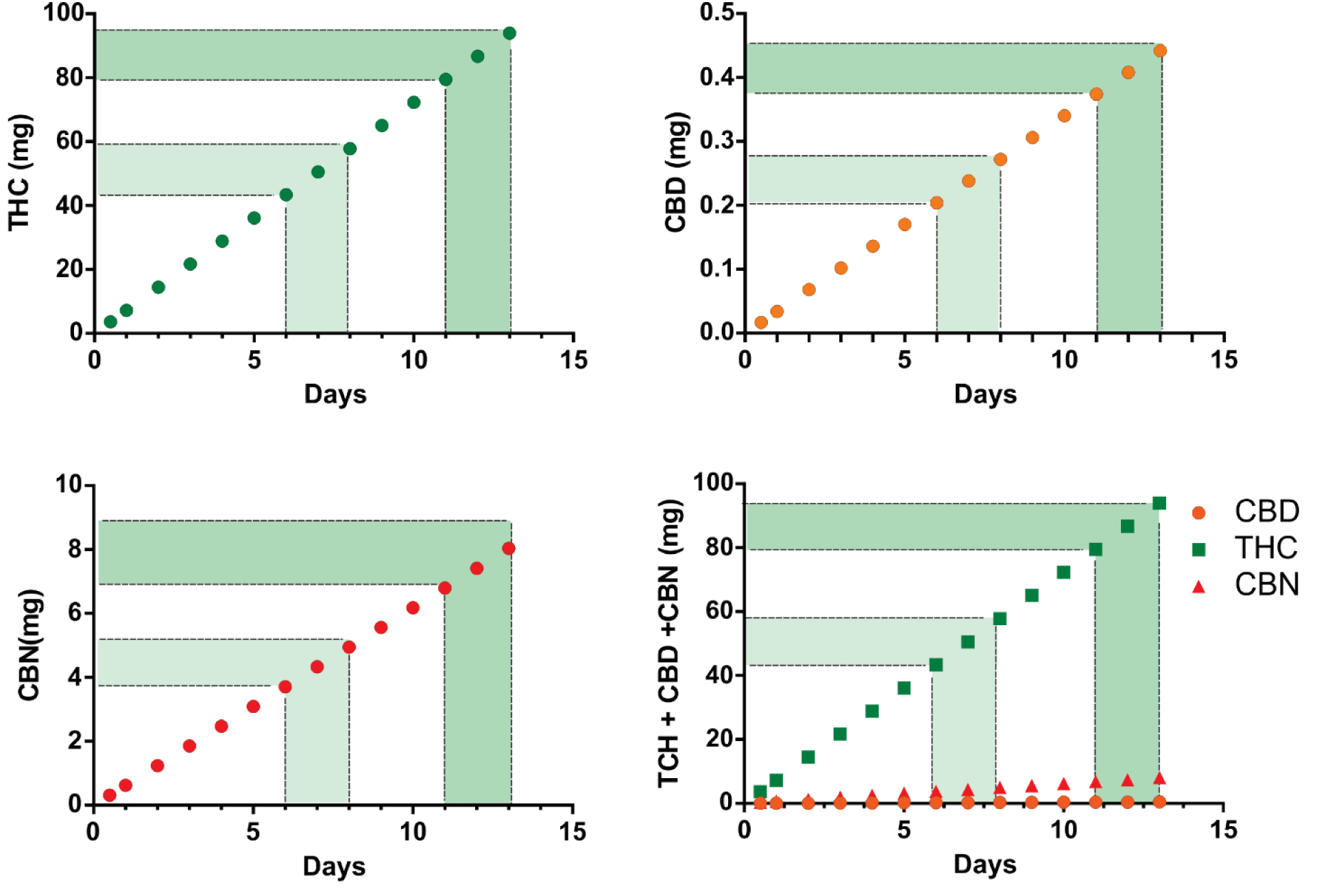
Final amount of principal cannabinoids provided during treatment. Rectangles indicate the accumulative amount of each component received by both groups. Group exposed between 6-8 days received (in mg): THC: 43.36-57.81, CBD: 0.204-0.272, CBN: 3.71-4.94. Group exposed during 11-13 days received (mg), THC: 79.48-93.94, CBD: 0.374-0.442, CBN: 6.79-8.03. Flies exposed five days before interruption received (in mg) THC: 36.13, CBD: 72.26, CBN: 6.18.

**Fig. S2.**
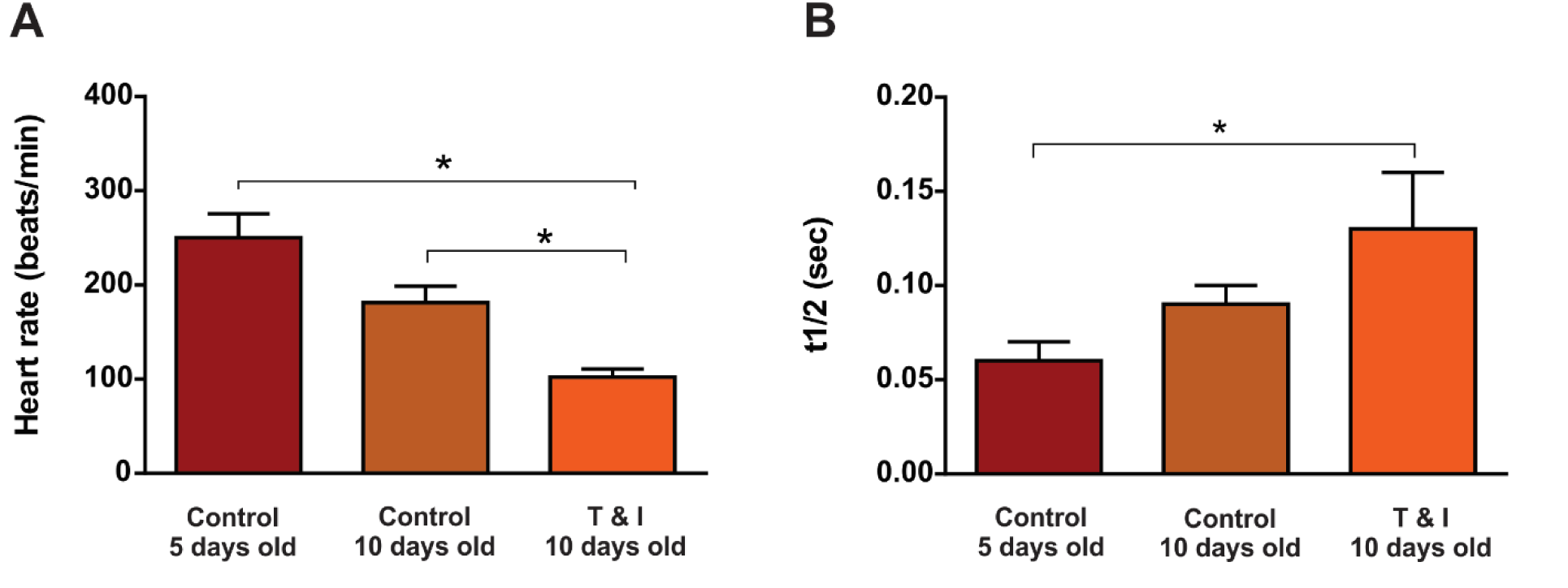
*Ca*nnabinoids are major contributors to heart rate reduction and t1/2 prolongation. Aging has a non-statistically incidence on both parameters when comparing 5 days old with 10 days old flies. 10 days old adult flies exposed to cannabinoids show a significant reduction in heart rate and an increase in t_1/2_ when compared to controls. Heart rate: 5 days old control flies: N = 14, 10 days old control flies N = 10, 10 days old adult flies (5 days and 5 days without cannabinoids, T&I) N = 11. t_1/2_: C 5d N = 14, C 10 d N = 10, T &I N = 11.

**Fig. S3.**
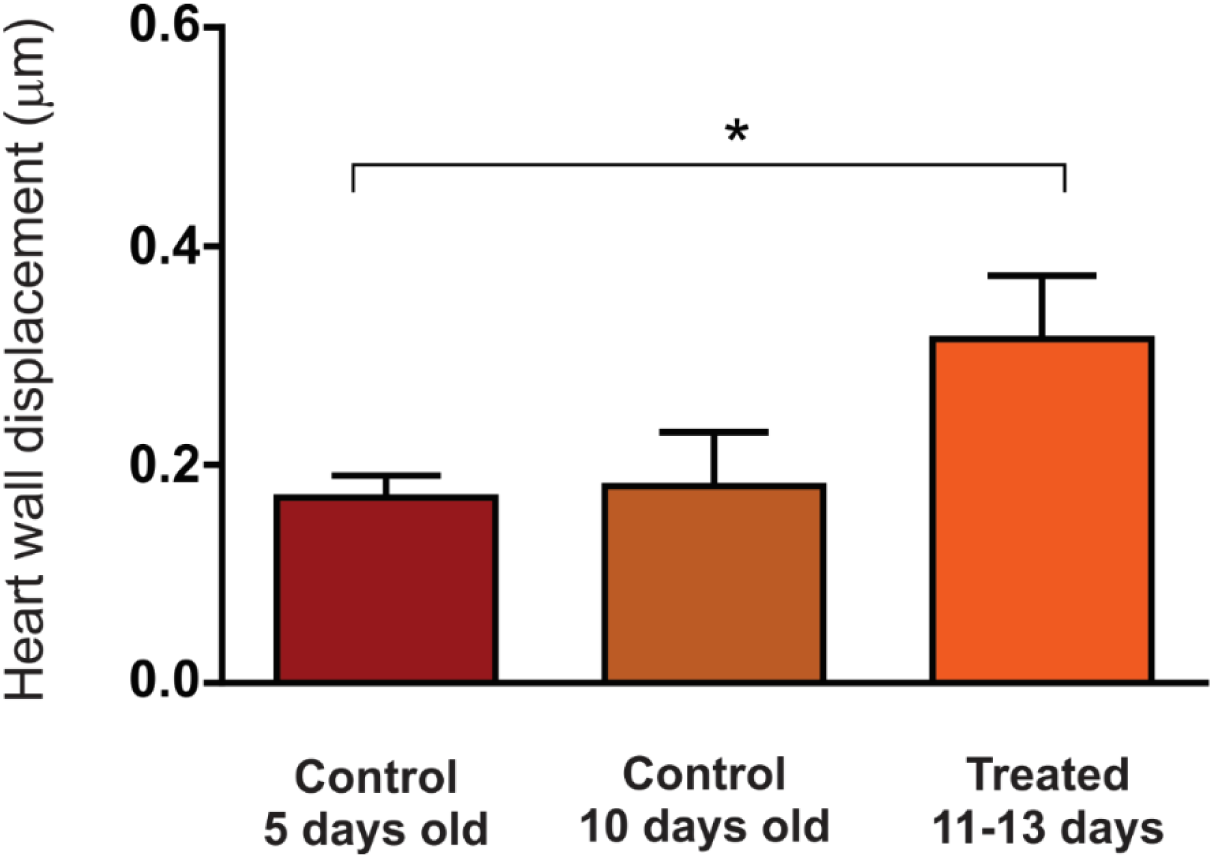
*Ca*nnabinoids are responsible of the increment in heart contractility. Non-treated 5 and 10 days old individuals exhibited similar contractility. The increment in this mechanical parameter in vaporized flies reflex the incidence of cannabinoids in this group. 5 days old control flies: N = 20, 10 days old control flies: N = 10, treated group: N = 7.

**Table S1.**
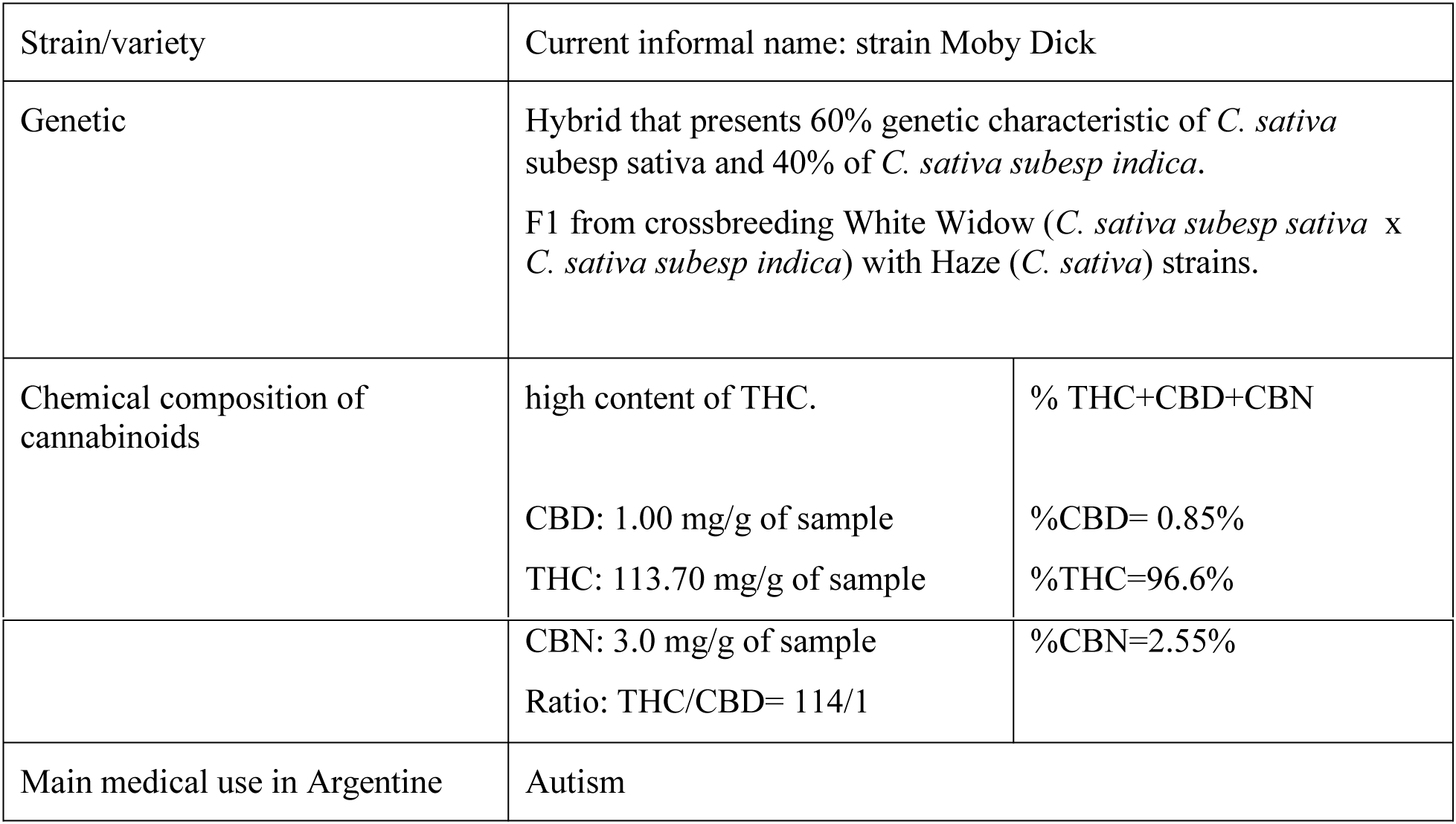
Genetic and chemical characterization of the studied strain of *Cannabis sp*. Vegetal material provided by the Argentinian NGO “Mamá Cultiva” was analyzed for determining % content of THC, CBD and CBN.

**Table S2.** Vaporized components. Temperature at which the main components present in *Cannabis sp* are vaporized in our conditions using a device setting at 188°C.

| **Cannabinoids** | **Boiling point (in °C)** |
| --- | --- |
| Delta 9- THC | 157 |
| CBD | 180 |
| CBN | 185 |
| **Terpenoids** |  |
| Beta myrcene | 166-168 |
| Beta caryophyllene | 119 |
| d-limonene | 177 |
| 1,8-cineole | 176 |
| Alpha-pinene | 156 |
| Beta-cymene | 177 |
| Delta-3-carene | 168 |
| **Flavonoids** |  |
| Apigenin | 178 |
| Cannflavin A | 182 |
| Beta-sitosterol | 134 |

